# Traumatic brain injury induced microglial and Caspase3 activation in the retina

**DOI:** 10.1101/2022.12.30.522345

**Authors:** Tamás Kovács-Öller, Renáta Zempléni, Boglárka Balogh, Gergely Szarka, Bálint Fazekas, Ádám J Tengölics, Krisztina Amrein, Endre Czeiter, István Hernádi, András Büki, Béla Völgyi

## Abstract

Traumatic brain injury (TBI) is among the main causes of sudden death after head trauma. These injuries can result in severe degeneration and neuronal cell death in the CNS, including the retina which is a crucial part of the brain responsible for perceiving and transmitting visual information. The long-term effects of mild-repetitive TBI (rmTBI) are far less studied thus far, even though damages induced by repetitive injuries occurring in the brain are more common, especially amongst athletes. rmTBI can also have a detrimental effect on the retina and the pathophysiology of these injuries are likely to differ from the severe TBI (sTBI) retinal injury.

Here we showed how rmTBI and sTBI can dissimilarly affect the retina. Our results indicate an increase in the number of activated microglial cells and Caspase3-positive cells in the retina in both traumatic models, suggesting a rise in the level of inflammation and cell death after TBI. The pattern of microglial activation appears evenly distributed and widespread but differs amongst the various retinal layers. sTBI induced microgial activation in both the superficial and deep retinal layers. In contrast to sTBI, no significant change occurred following the repetitive mild injury in the superficial layer, only the deep layer (spanning from the inner nuclear layer to the outer plexiform layer) shows microglial activation. This difference suggests that alternate response mechanisms play a role in the case of the different TBI incidents. The Caspase3 activation pattern showed a uniform increase in both the superficial and deep layers of the retina. This suggests a different action in the course of the disease in sTBI and rmTBI models and points to the need for new diagnostic procedures. Our present results suggest that the retina might serve as such a model of head injuries since the retinal tissue reacts to both forms of TBI and is the most accessible part of the human brain.

## 1. Introduction

The term Traumatic Brain Injury (TBI) is used as a collective term for pathological changes due to external forces that can lead to physiological, cognitive, and psychosocial disorders of the central nervous system. It is most commonly caused by accidents, sports injuries, and falls [1]. These injuries can result in a variety of disorders, disabilities, and sometimes seizures [2]. Road traffic accidents often lead to severe head injuries that can be effectively reduced by road measures. Moreover, there is an increased risk of head injuries among elderly patients, and athletes [3, 4]. Categorization of different types of head injury is challenging, as - mainly due to the diverse directions and magnitude of physical forces beside widely varying environmental conditions - the location, type, and extent of the induced pathoanatomical and pathophysiological sequelae may substantially differ case by case. [5].

The estimated rate of post-TBI and TBI-related deaths worldwide is more than 1.5 million per year [6] mostly as a result of the lack of diagnostic abilities and/or life-saving brain surgeries [7].

The intensity and speed of the different forces acting on the head determine the extent of tissue damage. The brain essentially moves in the cerebrospinal fluid and various harmful mechanical forces, including shear and rotation, induce the displacement of the brain and corresponding tissue damage [8]. Based on pathoanatomical considerations these lesions can be divided into focal and diffuse types. Focal damage usually occurs at the site of the impact or on the contralateral side of the brain. While the focal lesions (epidural-, subdural-, intracerebral hemorrhages and contusions) more often associated with moderate or severe head injuries in most of the TBI cases a mixture of focal and diffuse (edema, hypoxic-ischemic injury, microvascular injury, diffuse neuronal injury and diffuse axonal injury) types of pathological lesions develop (Mckee AC, Daneshvar DH. The neuropathology of traumatic brain injury. Handb Clin Neurol. 2015;127:45–66. (PMID: 25702209)). Diffuse pathological lesions - triggered by the shearing and rotational forces to the head - may occur in remote regions of the CNS (e.g. retina) compared to the direct impact site.

Based on the time course of the injury, we can speak of a primary injury that occurs at the time of the accident like fractures and intrusions. On the other hand, although secondary injury processes initiated at the moment of injury, become clinically apparent hours or days after the initial impact (edema, ischemia, hypoxia, neuroinflammatory processes) and may develop even years or decades after the injury (like chronic traumatic encephalopathy, post-traumatic stress disorder, dementia or endocrine deficiencies) causing severe quality of life issues for the affected patients as well as a huge socio-economic burden [9]. Therefore, there is an unmet clinical need for efficient diagnostic (and prognostic) possibilities. The retina is not only one of our most important sensory organs, but it is also a part of our CNS, directly linked to the brain and located peripherally thus providing easy access to diagnostic measures.

Studies show that characteristic retinal TBI symptoms include photophobia, double or blurred vision, visual paralysis, optic nerve disorders, and damaged or changed image processing (Fig. 1; [10]). TBI also includes retinal injury, which, according to current data, occurs in 20–40% of affected millions in TBI [11]. Mapping these lesions and also milder effects on the retina may help to understand the pathophysiology of TBI and may be an indicative factor in screening for severe cases. Our work is an important, pioneering research that seeks to expand our understanding of brain injuries and offers a potential diagnostic tool. Even when cellular and apoptotic lesions are present elsewhere in the brain, the retina can be a direct and most accessible indicator for TBI. However, at present, we lack the necessary information to connect direct retinal damage with TBI.

**Figure 1.**
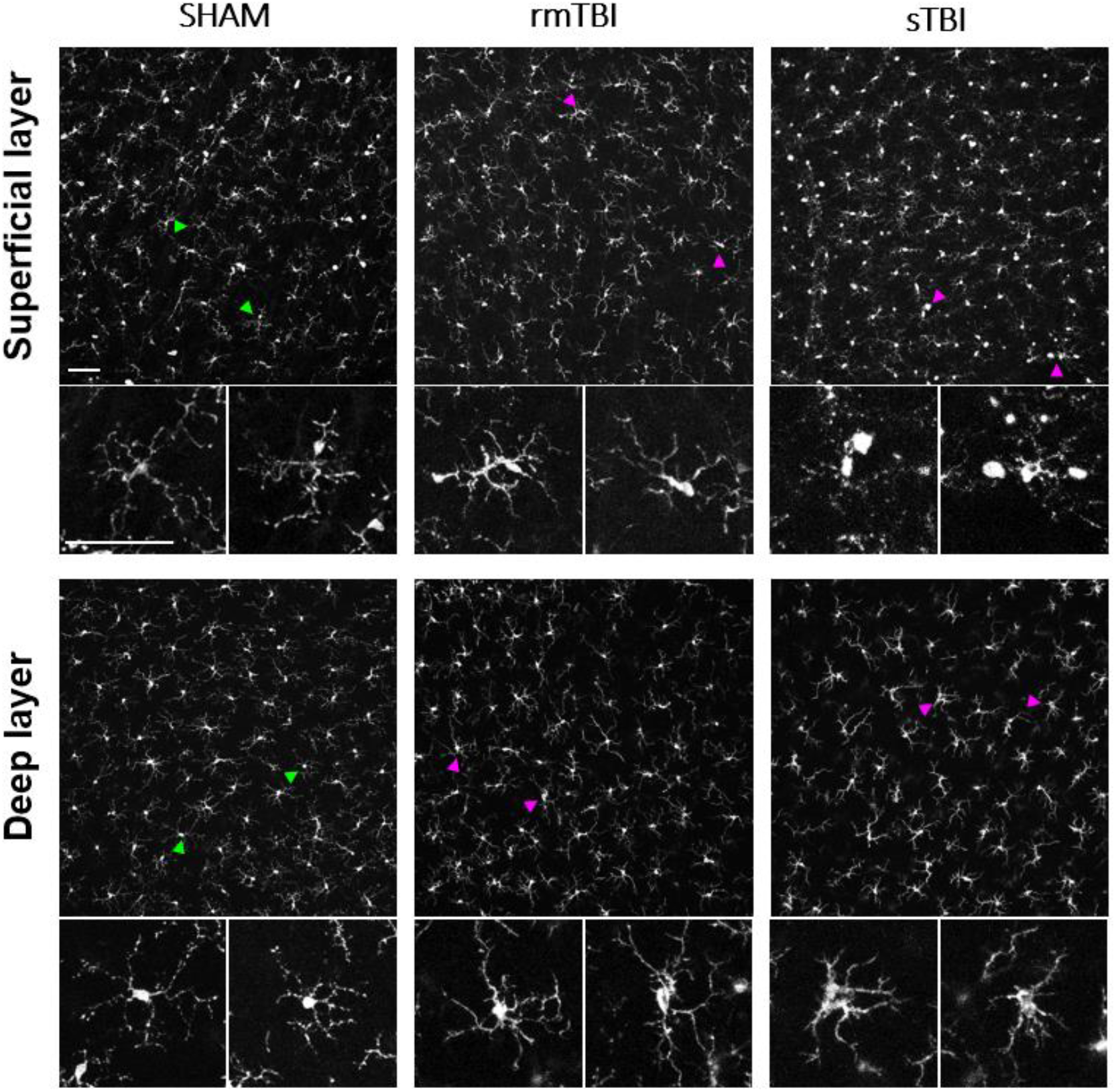
Microglial activation after sTBI and rmTBI in the retina. Microscope images show the activation of microglia sTBI (Marmarou 2 m) and rmTBI (3x Marmarou 15 cm) in the rat retina. Green arrowheads are showing examples of non-activated microglia from SHAM retinas, magnified on the bottom of each image. Magenta arrowheads showing microglia with activated morphology (described in Table 2) in rmTBI and sTBI. Scale Bar: 50 μm.

Microglia are the resident immune cells in the brain including the retina. They are immunocompetent cells constantly monitoring their environment to interact with possible threats. Microglial activation in the retina is present in a cohort of disease phenotypes [12, 13], hence it is important to understand how they contribute to the pathophysiology. In the central nervous system (CNS) their activation ranges on a continuum from neuroprotective to neurotoxic [14]. Their vast activation can contribute to further deterioration of the disease phenotype in the neuronal tissue via pro-inflammatory and phagocytosis mechanisms [12, 15]. Caspase3 (Casp3) is a central effector for apoptotic cell death. It can be widely activated in TBI [16], and retinal diseases or detrimental conditions [17]. Microglial activation is highly dependent on Caspase activation [18].

To the best of our knowledge, our study is the first to show how the retina is affected by sTBI and rmTBI. Latter is caused by multiple low-level impacts common among athletes (especially boxing, and soccer) and is increasingly recognized as a major cause of neurological diseases [19]. Our results show the rising numbers of activated microglial cells as well as activated Casp3 (act-Casp3) positive cells in the retina in both traumatic models. Our data reinforce views that emphasize the severity of rmTBI and suggest attention similar to sTBI.

Act-Casp 3 is a common executor of different apoptotic pathways induced by various damages including ischemia, excitotoxicity, and radiation [20]. Microglia can show elevated levels of act-Casp3 without cellular death as they have a bypass mechanism to avoid apoptosis [18].

This suggests a different action in the course of the disease and points forward to new diagnostic procedures for TBI.

## 2. Materials and Methods

### 2.1. Animals and preparation

Animal handling, housing, and experimental procedures were reviewed and approved by the ethical committee of the University of Pécs (BA02/2000-69/2017). Adult, male Long Evans rats (n=12, Charles River Laboratory, Germany) weighing 300-400 g were used in the experiment. All animals were treated in accordance with the ARVO Statement for the Use of Animals in Ophthalmic and Vision Research. All efforts were made to minimize pain and discomfort during the experiments.

To induce experimental TBI, we used an impact acceleration weight-drop model, originally published by Marmarou and Foda [21-24]. The steps of the surgical protocol are explained briefly as follows. Anesthesia was induced in an induction box with 5% isoflurane (Baxter, Hungary) in a 70:30 N2:O2 gas mixture. Once the anesthesia stabilized, we fixed the animal’s head in a stereotaxic frame. From this point, the anesthesia was carried out in 2 % isoflurane in the same gas mixture. After removing the hair from the animals’ scalp, we made a midline incision and removed the periosteum associated with the top of the skull. Halfway between the exposed bregma and lambda sutures, we fixed a stainless-steel disc, the so-called ‘helmet’ directly to the bone with cyanoacrylate glue. Then we laid the animal on a foam bed in a prone position. The helmet was positioned centrally under the weight-leading plexiglass tube. Experimental diffuse TBI was induced by dropping the weight from the height corresponding to the desired severity level. After TBI induction the helmet was removed and the surgical area was cleaned and disinfected. The wound was sutured and the animal was returned to its cage to recover. Through the surgical procedure, the physiologic parameters of the animals were monitored by a pulseoximeter (MouseOx Plus, Starr Life Corp., USA). Body temperature was monitored by Homotermic Monitoring System (Harvard Apparatus, USA) and maintained with a heating pad on the same device at 37 °C.

To investigate the acute pathological effects of experimental diffuse TBI on the retina we divided the animals into 3 groups (n=4 in each group). In the first, single severe TBI (sTBI) group we induced the injury with 450 g weight from a 2-meter height. In the second, repetitive mild TBI (rmTBI) group we used the same weight from a 15-centimeter height 5 times with 24 hours intervals. Finally, to eliminate the possible effects of anesthesia / environmental conditions, our third group (SHAM) did not receive weight-drop treatment, only anesthesia, and the surgical protocol of fixing the helmet on the skull. This experimental design allowed us to investigate pathological alterations between injuries of different degrees of severity and frequency.

Following the last treatments, the animals were sacrificed after a 24-hour survival. Control and TBI rats were perfused transcardially with 4% PFA (4% paraformaldehyde in PBS: 137 mM NaCl; 2.7 mM KCl; 10 mM Na2HPO4 * 7H2O, pH 7.4), and their eyes were immediately removed. The eyes were dissected in PBS by removing the cornea and lens. The resulting eyecups were additionally fixed in 4% PFA at room temperature for 15 minutes for better sample retention. After washing them three times for 10 minutes in PBS, the retinas were dissected from the eyecups, or the eyecups were kept for up to 3 weeks in 0.05% Na-azide in PBS at 4 ° C until processed.

### 2.2. Immunohistochemistry

Flat-mounted retinas were blocked in 100 μl of CTA (5% Chemiblocker, 0.5% TritonX-100, 0.05% Na-azide in PBS) overnight, room temp., humidified. After blocking, the retinas were treated with the primary antibodies (1000x + 1000x mouse SMI31 + SMI32 = SMI312, NE1022 / NE1023 - Calbiochem; 1000x rabbit Caspase3, AF835 - NovusBio; 2000x guinea pig Iba1, 234004 - SySy), diluted in CTA, for 48 h, room temperature (RT). Retinas were incubated for 48 hours at RT. After subsequent washing in PBS, three times, secondary antibodies were applied: 500x anti-rabbit Cy3 (715-165-150 - Jackson), 1000x anti-mouse A488 (A11017 - Invitrogen) and 1000x anti-mouse A647 (A21237 - Invitrogen) or anti-guinea pig A647 (A21450, Invitrogen) in CTA, and incubated overnight at room temperature (see Table 1 for abs.). After washing three times for 10 mins in PBS, they were coverslipped with Vectashield (Vector Laboratories, Peterborough, UK) using coverslip nr.1 (protocol based on and previously validated in [17, 25]).

**Table 1.** Antibodies

| Primary antibodies |  |  |  | Secondary antibodies, dyes |  |  |  |
| --- | --- | --- | --- | --- | --- | --- | --- |
| Name | Dilution | Source | Code | Name | Dilution | Source | Code |
| ms-SMI312 | 1:1000 | Calbiochem | NE1022 / NE1023 | anti-ms-Alexa488 | 1:1000 | Invitrogen | A11017 |
| gp-Iba1 | 1:2000 | SySy | 234004 | anti-ms-Alexa647 | 1:1000 | Invitrogen | A21237 |
| rb-Caspase-3 | 1:1000 | NovusBio | AF835 | anti-gp-Alexa647 | 1:1000 | Invitrogen | A21450 |
|  |  |  |  | anti-rb-Cy3 | 1:500 | Jackson | 715-165-150 |

### 2.3. Microscopy

Retinas were inspected using a Zeiss LSM 710 confocal laser scanning microscope (Plan Apochromat 10x, 20x, and 63x objectives (NA: 0.45, 0.8, 1.4, Carl Zeiss Inc., Jena, Germany) with normalized laser power and filter settings making 1.5 and 0.5 μm thin optical sections (further details in Balogh, 2021 [17]).

### 2.4. Measurement of Microglial and Casp3 activation

All measurements were performed using FIJI (NIH, USA, [26]). First, we performed two z-merges from the 5-5 optical stacks (3.75 μm) for the superficial and deep regions of MGs using only scans from mid-central retinal regions. The microglia were separated from the other signals based on their expression of Iba1 (*Ionized calcium-binding adapter molecule 1)*. Cells were manually grouped one by one according to their morphologies into activated and non-activated, using the ‘Cell-counter’ plugin in FIJI, according to the morphological classifications of Lawson and colleagues and others [27-29]. Only cells with the whole visible area were included, we omitted the ones on the edges.

We divided superficial (SL) and deep layers (DL) in the Z-stacks for MGs on the 20x images following the layers of blood vessels, overlapping NFL+GCL for SL and INL to ONL for DL. A ‘z-merge’ was done for these ranges from 5 to 5 consecutive optical sections. During microglial activation, counting and classifying cells with different morphologies (n = 1957 cells in 12 retinas) were performed one by one using the ‘cell counter’ plugin in FIJI. Differential morphological characteristics listed in Table 2 (based on the work of Davis and his colleagues [30]) were considered during selection. To preserve objectivity, the names of sample images were randomized.

**Table 2.** Morphological differences between non-activated and activated microglia.

| **Non-activated (naïve) morphology** | **Activated morphology** |
| --- | --- |
| Small, round soma | Enlarged, disorganized soma |
| Low soma to surround ratio | Soma to surroundings ratio increases |
| No amoeboid or leaf-like appendages are found | Amoeboid and leaf-like structures |
| Sporadic occurrence of act-MGs | Aggregate occurrence of act-MGs |

To examine Casp3 activation in the SL and DL we used the same z-merges, and the activated cells were separated from the background using other signals (GFAP, SMI312) together with the act-Casp3 signal to define retinal layer borders better and possible cell types affected.

### 2.5. Statistical analysis

Data was curated in MS Excel. One-way ANOVA analyses were performed using Origin18 (Origin, Version 2018b, OriginLab Corporation, Northampton, MA, USA.) and JASP (JASP Team (2022). JASP (Version 0.16.3)). Normal distribution was previously confirmed through statistical analysis.

## 3. Results

### 3.1 Effects of TBI on retinal microglia

In both rmTBI and sTBI samples massive microglial activation could be observed on merged stacks of the superficial layer (SL) and the deep layer (DL) (Fig 1). This activation was manifested in various morphological changes seen in Table 2. After counting and sorting microglial cells into activated/non-activated categories, according to the expressed morphological criteria (Table 2.), we found that TBI resulted in a significant increase in the number of activated microglia in both the deep layer (mean = 43.3; Figure 5; Table 2) and superficial MGs (mean = 64.35; Fig. 3; Table 3). However, this increase was observed only in sTBI among SL MGs, whereas only DL MGs were activated significantly in rmTBI (Fig 2.).

**Table 3.** Microglial activation by sTBI and rmTBI. Indicating the mean activated number and standard deviation (SD).

|  | SL-NA |  |  | SL-Act |  |  | SL-All |  |  |
| --- | --- | --- | --- | --- | --- | --- | --- | --- | --- |
|  | Sham | sTBI | rmTBI | Sham | sTBI | rmTBI | Sham | sTBI | rmTBI |
| AVG | 45.5 | 22.0 | 34.0 | 47.3 | 77.4 | 51.3 | 92.8 | 99.4 | 85.3 |
| SD | 17.6 | 20.0 | 1.7 | 18.3 | 18.9 | 13.6 | 18.4 | 6.8 | 15.0 |
|  | DL-NA |  |  | DL-Act |  |  | DL-All |  |  |
|  | Sham | sTBI | rmTBI | Sham | sTBI | rmTBI | Sham | sTBI | rmTBI |
| AVG | 62.5 | 21.2 | 26.0 | 10.8 | 46.6 | 41.0 | 73.3 | 67.8 | 67.0 |
| SD | 10.1 | 10.4 | 13.1 | 4.6 | 13.4 | 11.4 | 12.1 | 4.5 | 14.8 |

**Table 4.** Mean number of cells identified as act-Casp3 in TBI treatments in superficial retinal layers (NFL + GCL), indicating sample standard deviation (SD).

| SL | SHAM | sTBI | rmTBI |
| --- | --- | --- | --- |
| act-Casp3 cells | 15.50 | 165.20 | 117.75 |
| SD | 1.29 | 22.45 | 43.61 |

**Figure 2.**
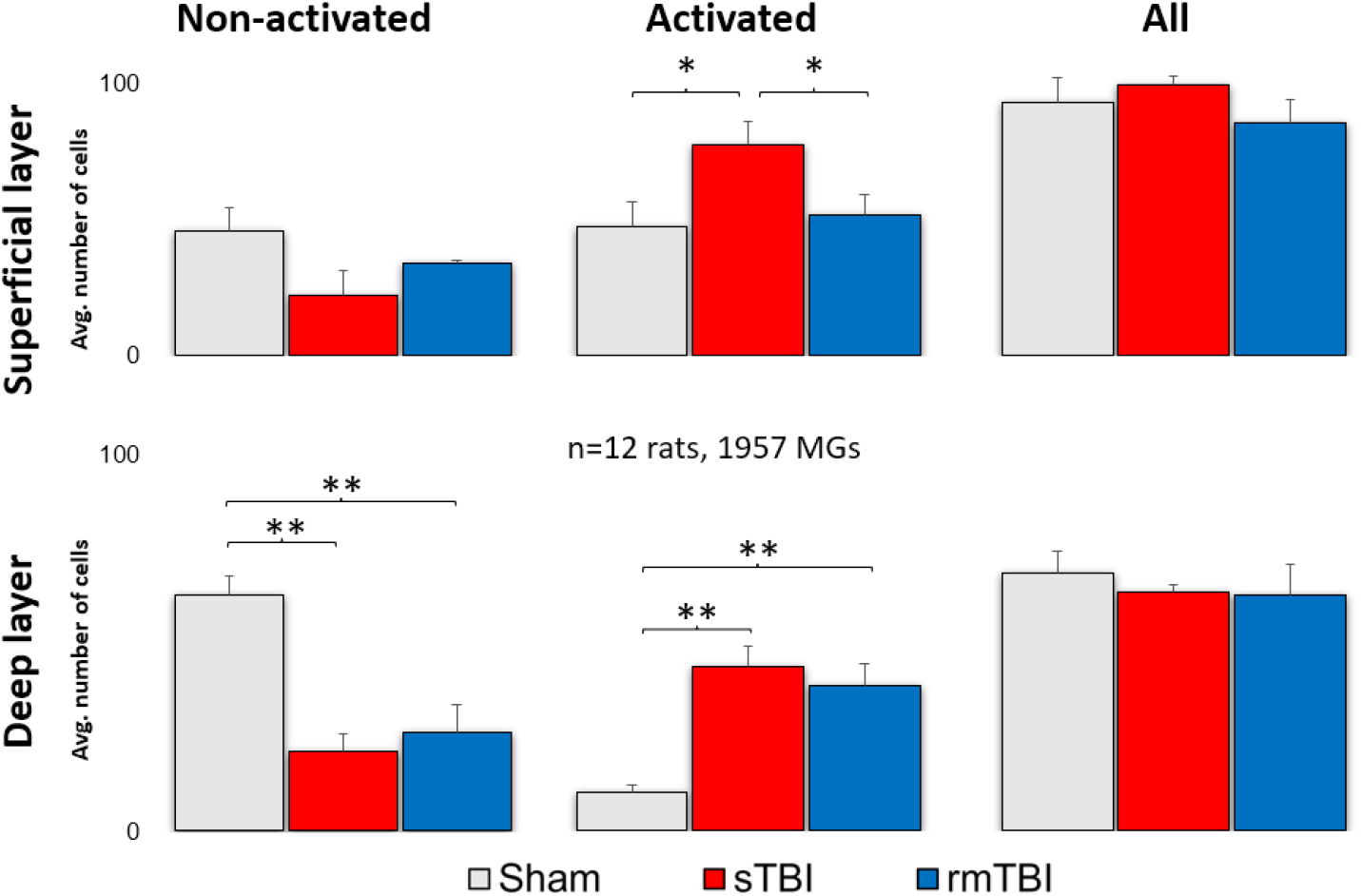
Microglial activation after sTBI and rmTBI in the retina. A significant increase in the number of activated microglia was observed after sTBI (Marmarou 2 m) and rmTBI (3x Marmarou 15 cm) in the deep layer of the rat retina. In the superficial layer, only sTBI resulted in microglial activation. There is no significant change in the total number of microglia. ANOVA, Gabriel post-hoc, * p <0.05, ** p <0.01.

**Figure 3.**
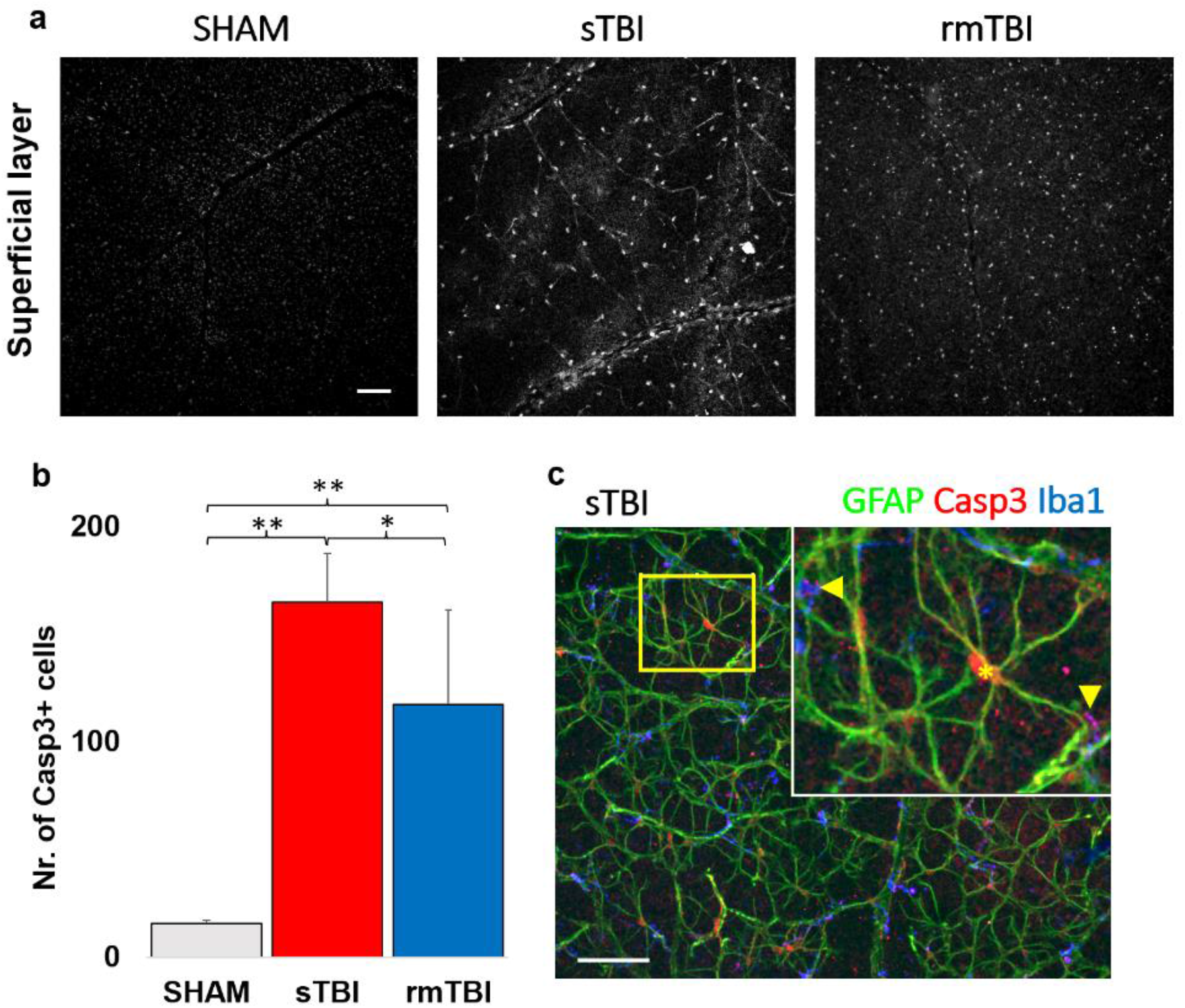
Casp3 activation in the SL of the retina. The figure shows an increase in the number of Casp3 + cells after sTBI and rmTBI compared to SHAM-operated animals (a). Bottom left (b) shows a plot of the mean increase in Casp3+ cell numbers (one-way ANOVA, Gabriel post-hoc, * p <0.05, ** p <0.01). (c) Following GFAP and Iba1 labeling, the Casp3+ cells were identified as astrocytes and microglia based on colocalizations with specific markers (microglia: arrows, astrocyte: *). Scale Bar: 50 μm.

### 3.2 Specific Caspase3 activation due to traumatic brain injury in the superficial layer of the retina

Casp3 activation was analyzed first in the superficial layers of the retina, where Casp3 activation was already observed in confocal LSM images of both sTBI and rmTBI samples, which unfolded in two ways. On one hand, it labeled the nuclei and the cytosol of cells of a distinct cell population in the neurofilament layer, and on the other hand, showed granular labeling in the ganglion cell layer (Fig. 3 c). We observed a large, 10.54x increase in the total number of act-Casp3 cells in sTBI, compared to SHAM (15.67 → 165.20), while a slightly smaller but still large 7.52x increase in activated cell number in rmTBI (15.67 → 117)., 75); (Figure 3b, Table 3). We should note that in the case of rmTBI, the standard deviation of our sample was much larger, but still showed a significant prominent increment (Fig. 3b, Table 3).

The large number of act-Casp3 cells described was identified using several markers (non-correlated markings are not provided here). Finally, Casp3 activation in SL was found in colocalizations with the two markers. These markers were GFAP (Glial-Fibrillar Acidic Protein) and Iba1 (Ionized calcium-binding adapter molecule 1e), the signaling of which is mainly restricted to astrocytes and microglia respectively in the retina [31, 32]. Using them, we identified these cells in SL as the main source of Casp3 + labeling (Fig. 3c).

Interestingly, the localization of the label differed between the two cell types. In GFAP+ astrocytes, the Casp3 signal was clearly restricted to the nucleus and its surrounding region, whereas the Casp3 localization of microglia was more limited to the protrusions. This could be observed in most of the cells in correlation with the Iba1 microglial marker (Fig. 3c, arrows).

### 3.2 Loss of axonal connections due to traumatic brain injury in the Neurofilament layer of the retina

In addition to the obvious microglial activation and Casp3 activation in the astrocytes in the SL of the rat retina SMI312 (Neurofilament H) labeling (prepared using SMI31 and 32 together) showed a decline in the axonal numbers in the NFL of the sTBI and rmTBI retinas in comparison to the SHAM (Fig4. a).

**Figure 4.**
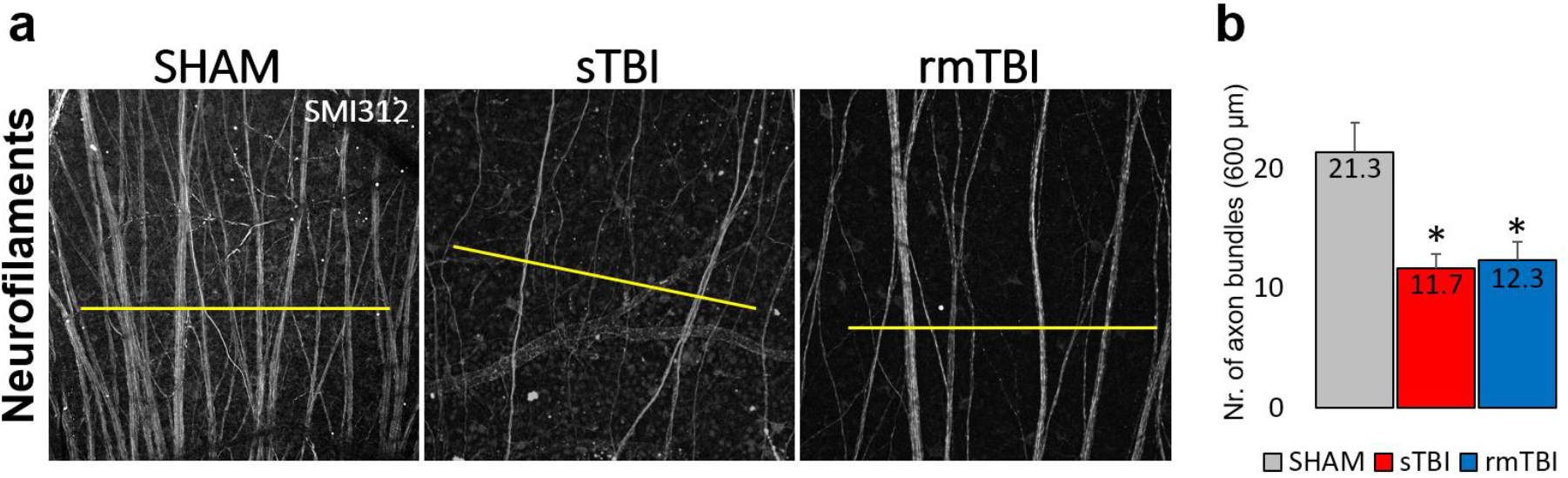
Change in SMI312 signal in the Neurofilament Layer in the retina after sTBI and rm TBI. (a) Images of the SMI312+ neurofilaments show a decline in both sTBI and rmTBI. 600 μm lines (yellow) were used to measure peaks of axon bundles after plot profile analysis, whereas the average number is shown by (b) indicating the crossing axon bundles (n=3 each). Paired t-test, * p <0.05.

Profile plot analysis of the neuronal marker SMI312 revealed that the positive axon bundles declined in both sTBI (-45.3%) and rmTBI (-42.2%) if measured from the same mid-central retinal regions (Figure 4). This indicates an early onset of neuronal decline in TBI having similar effects from a low level of multiple impacts and one severe impact.

### 3. Caspase3 is activated due to traumatic brain injury in the DL of the retina

After closer examination we identified the Casp3+ cells mainly as bipolar cells based on their soma diameters (mainly between 4-7 um) [33] and clear co-labeling with NT. A second cohort of cells appeared non-NT-labeled with bright Casp3 labeling. The soma diameters of these latter cells were larger compared to bipolar cells and the somas were located distally from the innermost sublamina of the INL where amacrine cells can be observed (based on the strong NT labels and the relatively large, 7-14 μm somata) [34]). Therefore, this second cohort of non-NT-labeled Casp3 + cells were identified as Müller cells.

Subsequent analyses of the Casp3 labeling in the DL unfolded in two ways. A massive (21.5 and 39.9 fold) increase in Casp3 activation was obvious after counting the individual cells with the help of NT labeling (Fig5 b,c, Table 5) in both the rm- and sTBI samples.

**Figure 5.**
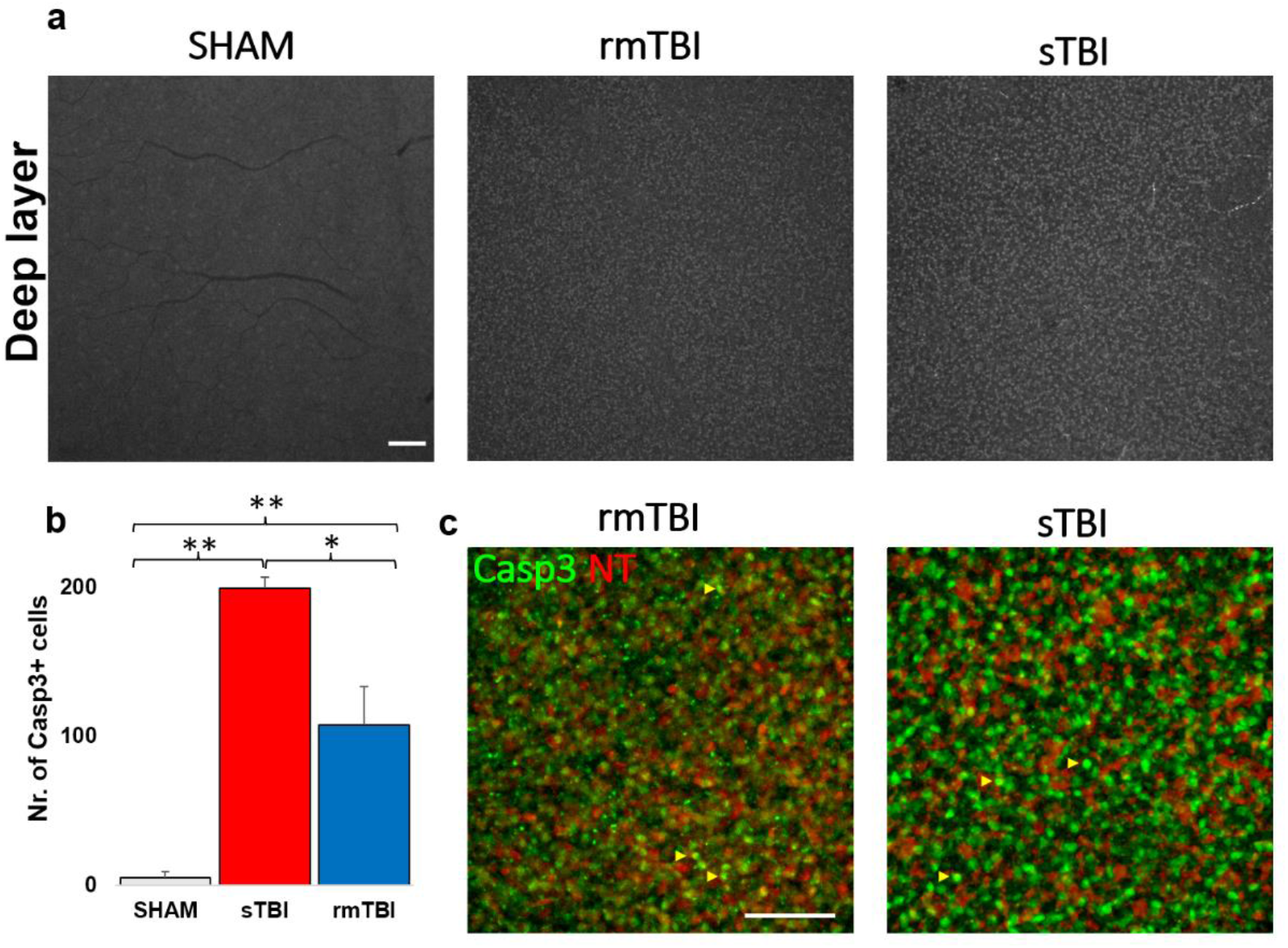
Casp3 activation in the deep layers of the retina. The figure shows an increase in the number of Casp3 + cells (a-gray, b-green) after sTBI and rmTBI compared to SHAM-operated animals (a). (b) shows a plot of the mean increase in Casp3 + cell numbers (one-way ANOVA, Gabriel post-hoc, * p <0.05, ** p <0.01; c) Neurotrace-640 (NT, red) labeling helped to identify these cells as bipolar cells and possibly Müller cells on the magnified images (‘c’, shown with yellow arrows, not labeled with NT; Scale Bar: 50 μm).

**Table 5.**
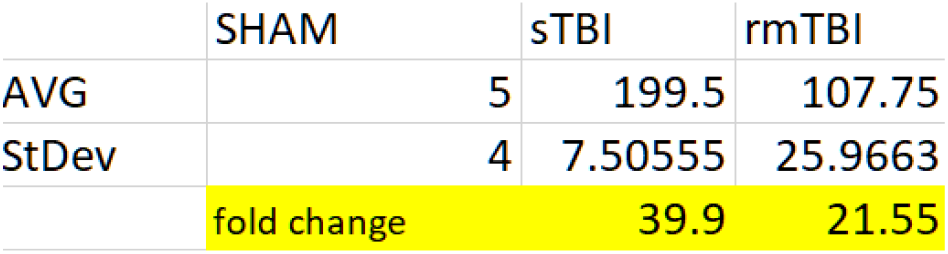
Average act-Casp3 cell counts in TBI in the deep layers of the retina.

## Discussion

TBI pathophysiology remains poorly understood. It results from direct impact or acceleration-deceleration to the head. The details and markers are still widely being explored. The retina is not only part of the CNS but it is directly connected to other parts of the brain, since the retina’s output cells, the Retinal Ganglion Cells (RGCs), actively project their axons to over 40 subcortical brain regions including image-forming (dorsal lateral geniculate nucleus) and non-image forming pathways (superior colliculus, medial terminal nucleus, nucleus of the optic tract, basal medial preoptic area, ventrolateral preoptic area, suprachiasmatic nucleus, medial pedunculopontine nucleus, etc.) [35, 36]. These RGC axons are openly exposed to the effects of different injuries (sTBI or rmTBI), therefore, the retina can actively intercept signals from the brain tissue, projected by axonal connections and indicated by the cells in the retina (Lukas, T.J., Wang, A.L., Yuan, M. et al. Early cellular signaling responses to axonal injury. Cell Commun Signal 7, 5 (2009). https://doi.org/10.1186/1478-811X-7-5).

### 1. Microglial activation due to traumatic brain injury

Based on our results, microglial activation clearly occurred in both the sTBI and rmTBI models. This suggests that the assumption from previous research about only 40% of TBI patients having a negative retinal impact appears to be highly underestimated [11]. Surprisingly, although we observed a high level of activation at sTBI, the data measured in rmTBI was not far behind those measured in sTBI, which may be alarming in cases where someone (e.g., sports such as football, boxing) is continuously exposed to minor cranial injuries [37].

Microglial cells continuously scan their environment, and their activation state faithfully reflects the state of the retina and that changes are taking place in the tissue, hard-wired to diverse parts of the brain [38]. The activation status of microglia may be different (M0,-1,-2) and treatment options may vary accordingly. Many drugs (e.g. CB2 inverse agonists) can affect the activation state of microglia and can reverse the noxious inflammatory M1 phase to the reparative M2 [39]. The differential activation of microglia can be of further interest for the right treatment of TBI-induced disease.

However, in addition to the choice of treatment options, activation of microglial cells could be used as biomarkers of TBI in the future, as these cells may be visualized by a combination of different staining procedures and fluorescent coherent optical tomography (fOCT) methods. As not all lesions can be traced with current imaging methods, the development of new in vivo procedures is needed. However, no real breakthrough has yet been made in the development of imaging methods in this area [40].

Activated microglia can release a number of factors (e.g., tumor necrosis factor ∝, interleukin-1β) that can induce inflammatory and degenerative processes in the retina [41]. On the other hand, the neuroprotective effect of microglia is also known, which can be a major help in preserving vision [42]. The detection and further examination of these factors represent great advances in determining the exact role of microglia in TBI.

### 2. Caspase3 activation, cell death marker

Caspases belong to the family of cysteine proteases. In mammals, 14 different caspases have been discovered that play a role in both apoptosis and inflammatory processes [17, 43, 44]. Animal studies have demonstrated activation of caspases (3, 7, 9, 12) in TBI in the brain [45, 46], where Casp3 activation is a key, cumulative marker of cell death since it signals the onset of both the intrinsic and extrinsic pathways of apoptosis, as it connects the two pathways. Because unregulated cell death would be detrimental, caspases are stored as latent zymogens in the cells, where they are converted to active caspases by proteolytic cleavage. Earlier studies suggest that not only is this cleavage required for activation, but also for a dimerization that binds inactive monomers together [47, 48]. We chose Casp3, together with other markers, to summarize apoptotic processes. According to our results, the SL clearly indicates Casp3 activation in GFAP+ cells, which we identified as astrocytes. Astrocyte activation also occurs in high Intra Ocular Pressure (IOP) conditions, such as glaucoma [49].

Further markers can subserve to identify apoptotic cell types. However, in the DL of the retina, it is difficult to identify distinct cell types expressing the Casp3+ signal due to the greater number of cells, the many cell types the relatively small soma sizes as well as the lack of cell type-specific markers (e.g. amacrine, bipolar subtypes). On the other hand, various cell types involved can be more easily identified with cell-type-specific markers labeling the SL cells with fine morphological detail, therefore we used SMI312 and GFAP. By quantifying the labeling, and also adding SMI312 and GFAP, resulting in clear cellular morphology, we were able to identify the activated cells. In the GFAP+ population, the cells’ nuclei were co-labeled with Casp3, and we were able to visualize subcellular granular Casp3+ labeling colocalized with Iba1. As GFAP labels the astrocytes, and Iba1 the MGs in the retina [50, 51], we identified them as the most affected population in both TBI models. However, the differential labels in the two cell populations may indicate that Casp3 might have different effects on them.

The neuronal marker SMI312, in some cells, coexpressed with Casp3 indicates the involvement of RGCs. However, we did not encounter extensive labeling of neurons in the examined samples from either rm-or sTBI, the loss of immunopostive axons also show the same effect on RGCs (measured on n=3). It should be noted that the Casp3+ signal did not indicate that a large number of cells had been committed toward cell death, however, we did not perform extensive colocalization studies in this study. The axonal degeneration is obvious by the lack of SMI312+ axonal labeling in both sTBI and rmTBI. These are homogeneously affected by the axonal labeling in the retinas. This result thus coincided with neuronal survival in similar works by other research groups [52]. However, it may be that more neuronal Casp3 activation could have been observed with longer survival time, giving more time for apoptotic processes.

The extent of Diffuse Axonal Injury (DAI) in sTBI and rmTBI may come from other brain areas. This type of injury first arises in the axonal head and by disorganization of the cytoskeleton and transport mechanisms, this in turn deteriorates the axon and the soma of the affected cell through a Ca2+ and calpain-induced process [53]. Even now, it is still a matter of debate if the surviving GCs will be able to maintain normal visual function in a long term [52, 54, 55], thus further studies are required in the future in this subject.

In addition to neuronal colocalization, the use of GFAP was evident in the retina, as activation of astrocytes by TBI could also be expected. The results we obtained show that astrocytes began large-scale mass activation of Casp3, providing a clear sign of apoptosis. The astrocyte network is responsible for the health of retinal neurons by maintaining homeostasis and involvement in the blood-retinal barrier [56]. Injury to the eye and retina can trigger more secondary processes than neogliosis, which is characterized by elevated levels of intermediate filaments of glial cells, including GFAP, vimentin, and nestin. Furthermore, microdomains of astrocyte reactivity can be biomarkers of functional decline of RGCs, so diagnostic imaging of astrocytes in the nerve fiber layer using modern imaging technologies can be an effective solution to detect axonal transport deficiency that can occur in such lesions (and, for example, glaucoma).

We observed Casp3 activation of microglia in both rm- and sTBI. However, it is important to note that activation of Casp3 in microglia does not necessarily imply a commitment toward cell death, as these cells have a bypass mechanism to avoid cell death. In microglial cells, various inflammatory factors induce the activation of Casp8 and then Casp3. Active Casp3, in turn, promotes the activation of microglial inflammatory pathways through a protein kinase Cδ-dependent pathway without induction of cell death [18]. Activation of Casp3 occurs as a two-step process in which the zymogen is first cleaved by upstream caspases, such as Casp8, to form an intermediate but active cytoplasmic p19 / p12 complex. An autocatalytic process then creates the fully mature form of the enzyme p17 / p12, which is then transferred to the nucleus. Induction of the cellular inhibitor of apoptosis protein 2 (cIAP2) upon microglial activation prevents the conversion of the p19 subunit of Casp3 to the p17 subunit, which is responsible for the cessation of Casp3 activity. By reversing the cIAP2-dependent process, the repressive effects are exerted, reactivate the inflammatory function of microglial cells, and may eventually promote their death [57]. We can assume that somehow Casp3 accumulation might act as a key for phenotype change in the end.

Considering this, however, in our case we do not expect extensive cell death in the high number of Casp3 + microglia. Instead, our results, as such, are further evidence of the activation of microglial cells beyond the morphological features. These microglia do not necessarily induce cell death and phagocyte, but may be involved in tissue protection [58] for cells in which Casp3 activation is not associated with the process. Therefore, Casp3 activation can be an additional marker for TBI.

### 3. Fate of different cell types under the influence of TBI

As mentioned in combination with Casp3, other cell-specific markers can be identified. However, with the markers we used, we could only draw the following conclusions. Only a limited number of markers could be used in the experiments. Unfortunately, we could not co-administer SMI312 with Casp3 in the entire sample set due to the lack of available secondary channels in IHC. However, no cell-specific Casp3 activation was examined with further specific markers, but damage to other cell types could not be completely ruled out. This can be observed through further experiments in the future.

GFAP may be expressed in Müller cell endings [59] in some cases due to severe damage, however, no Müller cell endings were observed on the GFAP label morphologically based on our experiments. Since no Müller cell-specific marker was used (e.g., glutamine synthetase, Sox9, RLBP9; [60], their involvement cannot be completely ruled out based on our study.

SMI312 labels both the non-phosphorylated and phosphorylated neurofilament-H (NF-H) and are therefore specific for RGCs [61]. This marker potentially labels living cells, hence the intermediate filaments decompose in apoptosis [62]. NF-H can also be detected in serum after the onset of DAI peaking at 12-48 h after injury [63, 64]. NF-H is considered the most convenient marker of DAI diagnosis and together with MG-imaging, this might be the next step for a more accurate diagnosis of TBI pathology in the future.

In conclusion, we showed that both sTBI and rmTBI comparably had a detrimental effect on the retina with the exception of DL activation, where rmTB had less effect. We showed that act-Casp3 is elevated in numerous cells, identified astrocytes, and microglia as major contributors. In the DL act-Casp3 is present in bipolars and Müller cells. We also identified a progressive decline in SMI312+ axonal bundles suggesting an interference with further cellular connections. Our results highlight the importance of retinal involvement in TBI and suggest that monitoring the retina’s health status could be utilized as a future biomarker tool for diagnostic purposes, able to detect the fine changes shown here.

## Supplementary Materials

### Author Contributions

conceptualization, T.KÖ.; methodology, T.KÖ., R.Z., B.F., K.A., B.B., G.S.; validation, T.KÖ., B.B, G.S., B.V.; formal analysis, T.KÖ., R.Z., B.B., G.S., A.J.T.; investigation, T.KÖ., R.Z., B.B., G.S.; resources, B.V., A.B., I.H.; data curation, T.KÖ., G.Sz., R.Z. A.J.T., B.B.; writing—original draft preparation, T.KÖ., R.Z., B.B., G.S.; writing—review and editing, B.V., T.KÖ., G.S., E.C., B.F., A.J.T.; visualization, T.KÖ., R.Z.; supervision, T.KÖ., B.V., A.B., E.C.; funding acquisition, B.V., A.B., E.C., I.H., B.B. G.S.; see [CRediT taxonomy](https://img.mdpi.org/data/contributor-role-instruction.pdf)

### Funding

This study was supportedby the European Union under the action of the ERA-NET COFUND (2019-2.1.7-ERANET-2021-00018) to B.V., by the Hungarian Brain Research Program 2 (2017-1.2.1.-NKP-2017) to B.V., by NKFIH (OTKA NN128293) to B.V. as well as (OTKA K134555) to A.B., E.C. and K.A., by the European Union and the State of Hungary, co-financed by the European Social Fund in the framework of TAMOP-4.2.4.A/ 2-11/1-2012-0001 National Excellence Program (B.V.); by the ÚNKP-20-1-I-PTE-801 New National Excellence Program of the Ministry for Innovation and Technology (to BB); by the ÚNKP-22-3-II-PTE-1414 New National Excellence Program of the Ministry for Innovation and Technology (to G.S.). Project no. TKP2021-EGA-16 has been implemented with the support provided from the National Research, Development and Innovation Fund of Hungary, financed under the TKP2021-EGA funding scheme. The research was performed in collaboration with the Histology and Light Microscopy core facility at the Szentágothai Research Centre of the University of Pécs with support from GINOP-2.3.2-15-2016-00036.

## Acknowledgment

We acknowledge the administrative help provided by the Szentágothai Research Centre and technical support of the Histology and Light Microscopy and Nano-Bio-Imaging core facilities located at the research center. We thank the staff of the Animal Core facility of the SZRC, University of Pécs.

## Conflicts of Interest

The authors declare no conflict of interest.

## Notes

### Competing Interest Statement

The authors have declared no competing interest.

## References

1. Orff, H.J.; Ayalon, L.; Drummond, S.P. Traumatic brain injury and sleep disturbance: a review of current research. J Head Trauma Rehabil. 2009, 24(3), 155–65. doi: 10.1097/HTR.0b013e3181a0b281.

2. Jennekens, N.; de Casterlé, B.D.; Dobbels, F. A systematic review of care needs of people with traumatic brain injury (TBI) on a cognitive, emotional and behavioural level. J Clin Nurs. 2010, 19(9-10), 1198–206. doi: 10.1111/j.1365-2702.2009.03114.x.

3. Harvey, L.A.; Close, J.C. Traumatic brain injury in older adults: characteristics, causes and consequences. Injury. 2012, 43(11), 1821–6. doi: 10.1016/j.injury.2012.07.188.

4. Ling, H.; Hardy, J.; Zetterberg, H. Neurological consequences of traumatic brain injuries in sports. Mol Cell Neurosci. 2015 66(Pt B):114–22. doi: 10.1016/j.mcn.2015.03.012.

5. Hamel, R.N.; Smoliga, J.M. Physical Activity Intolerance and Cardiorespiratory Dysfunction in Patients with Moderate-to-Severe Traumatic Brain Injury. Sports Med. 2019, 49(8), 1183–1198. doi: 10.1007/s40279-019-01122-9.

6. DE Souza, R.L.; Thais, M.E.; Cavallazzi, G.; Paim Diaz, A.; Schwarzbold, M.L.; Nau, A.L.; Rodrigues, G.M.; Souza, D.S.; Hohl, A.; Walz, R. Side of pupillary mydriasis predicts the cognitive prognosis in patients with severe traumatic brain injury. Acta Anaesthesiol Scand. 2015, 59(3), 392–405. doi: 10.1111/aas.12447.

7. van Dijck, J.T.J.M.; Bartels, R.H.M.A.; Lavrijsen, J.C.M.; Ribbers, G.M.; Kompanje, E.J.O.; Peul, W.C. The patient with severe traumatic brain injury: clinical decision-making: the first 60 min and beyond. Curr Opin Crit Care. 2019, 25(6), 622–629. doi: 10.1097/MCC.0000000000000671.

8. Zhou, Z.; Li, X.; Kleiven, S. Biomechanics of Periventricular Injury. J Neurotrauma. 2020, 15;37(8), 1074–1090. doi: 10.1089/neu.2019.6634.

9. Davis, A.E. Mechanisms of traumatic brain injury: biomechanical, structural and cellular considerations. Crit Care Nurs Q. 2000, 23(3), 1–13. doi: 10.1097/00002727-200011000-00002.

10. Das, M.; Tang, X.; Mohapatra, S.S.; Mohapatra, S. Vision impairment after traumatic brain injury: present knowledge and future directions. Rev Neurosci. 2019, 30(3), 305–315. doi: 10.1515/revneuro-2018-0015.

11. Bodnar, C.N.; Watson, J.B.; Higgins, E.K.; Quan, N.; Bachstetter, A.D. Inflammatory Regulation of CNS Barriers After Traumatic Brain Injury: A Tale Directed by Interleukin-1. Front Immunol. 2021 12, 688254. doi: 10.3389/fimmu.2021.688254.

12. Guo, L.; Choi, S.; Bikkannavar, P.; Cordeiro, M.F. Microglia: Key Players in Retinal Ageing and Neurodegeneration. Front Cell Neurosci. 2022, 16, 804782. doi: 10.3389/fncel.2022.804782.

13. Rashid, K.; Akhtar-Schaefer, I.; Langmann, T. Microglia in Retinal Degeneration. Front Immunol. 2019 10, 1975. doi: 10.3389/fimmu.2019.01975.

14. Hellwig, S.; Heinrich, A.; Biber, K. The brain’s best friend: microglial neurotoxicity revisited. Front Cell Neurosci. 2013 7, 71. doi: 10.3389/fncel.2013.00071.

15. Shi, Y.; Manis, M.; Long, J.; Wang, K.; Sullivan, P.M.; Remolina Serrano J, Hoyle R, Holtzman DM. Microglia drive APOE-dependent neurodegeneration in a tauopathy mouse model. J Exp Med. 2019 216(11), 2546–2561. doi: 10.1084/jem.20190980.

16. Clark, R.S.; Kochanek, P.M.; Watkins, S.C.; Chen, M.; Dixon, C.E.; Seidberg, N.A.; Melick, J.; Loeffert, J.E.; Nathaniel, P.D.; Jin, K.L.; Graham, S.H. Caspase-3 mediated neuronal death after traumatic brain injury in rats. J Neurochem. 2000 74(2), 740–53. doi: 10.1046/j.1471-4159.2000.740740.x.

17. Balogh, B.; Szarka, G.; Tengölics, Á.J.; Hoffmann, G.; Völgyi, B.; Kovács-Öller, T. LED-Induced Microglial Activation and Rise in Caspase3 Suggest a Reorganization in the Retina. Int J Mol Sci. 2021, 22(19), 10418. doi: 10.3390/ijms221910418.

18. Burguillos, M.A.; Deierborg, T.; Kavanagh, E.; Persson, A.; Hajji, N.; Garcia-Quintanilla, A.; Cano, J.; Brundin, P.; Englund, E.; Venero, J.L.; Joseph, B. Caspase signalling controls microglia activation and neurotoxicity. Nature. 2011, 472(7343), 319–24. doi: 10.1038/nature09788.

19. Eyolfson, E.; Khan, A.; Mychasiuk, R.; Lohman, A.W. Microglia dynamics in adolescent traumatic brain injury. J Neuroinflammation. 2020, 17(1), 326. doi: 10.1186/s12974-020-01994-z.

20. McIlwain, D.R.; Berger, T.; Mak, T.W. Caspase functions in cell death and disease. Cold Spring Harb Perspect Biol. 2013, 5(4), a008656. doi: 10.1101/cshperspect.a008656.

21. Marmarou, A.; Foda, M.A.; van den Brink, W.; Campbell, J.; Kita, H.; Demetriadou, K. A new model of diffuse brain injury in rats. Part I: Pathophysiology and biomechanics. J Neurosurg. 1994, 80(2), 291–300. doi: 10.3171/jns.1994.80.2.0291.

22. Chakraborty, N.; Hammamieh, R.; Gautam, A.; Miller, S.A.; Condlin. M.L.; Jett, M.; Scrimgeour, A.G. TBI weight-drop model with variable impact heights differentially perturbs hippocampus-cerebellum specific transcriptomic profile. Exp Neurol. 2021, 335, 113516. doi: 10.1016/j.expneurol.2020.113516.

23. Johnson, V.E.; Meaney, D.F.; Cullen, D.K.; Smith, D.H. Animal models of traumatic brain injury. Handb Clin Neurol. 2015, 127, 115–28. doi: 10.1016/B978-0-444-52892-6.00008-8.

24. Büchele, F.; Morawska, M.M.; Schreglmann, S.R.; Penner, M.; Muser, M.; Baumann, C.R.; Noain, D. Novel Rat Model of Weight Drop-Induced Closed Diffuse Traumatic Brain Injury Compatible with Electrophysiological Recordings of Vigilance States. J Neurotrauma. 2016, 33(13), 1171–80. doi: 10.1089/neu.2015.4001.

25. Kovács-Öller, T.; Szarka, G.; Tengölics, Á.J.; Ganczer, A.; Balogh, B.; Szabó-Meleg, E.; Nyitrai, M.; Völgyi, B. Spatial Expression Pattern of the Major Ca2+-Buffer Proteins in Mouse Retinal Ganglion Cells. Cells. 2020, 9(4), 792. doi: 10.3390/cells9040792.

26. Schindelin, J.; Arganda-Carreras, I.; Frise, E.; Kaynig, V.; Longair, M.; Pietzsch, T.; Preibisch, S.; Rueden, C.; Saalfeld, S.; Schmid, B.; et al. Fiji: an open-source platform for biological-image analysis. Nature methods 2012, 9(7), 676–682. doi: 10.1038/nmeth.2019

27. Lawson, L.J.; Perry, V.H.; Dri, P.; Gordon, S. Heterogeneity in the distribution and morphology of microglia in the normal adult mouse brain. Neuroscience 1990, 39(1), 151–170. doi: 10.1016/0306-4522(90)90229-w

28. Davis, E. -J.; Foster, T. D.; Thomas, W. E. Cellular forms and functions of brain microglia. Brain research bulletin 1994, 34(1), 73–78. doi: 10.1016/0361-9230(94)90189-9

29. Streit, W. J.; Walter, S. A.; Pennell, N. A. Reactive microgliosis. Progress in neurobiology 1999, 57(6), 563–581. doi: 10.1016/s0301-0082(98)00069-0

30. Davis, B.M.; Salinas-Navarro, M.; Cordeiro, M.F.; Moons, L.; De Groef, L. Characterizing microglia activation: a spatial statistics approach to maximize information extraction. Sci Rep. 2017, 7(1), 1576. doi: 10.1038/s41598-017-01747-8.

31. Schnitzer, J. Distribution and immunoreactivity of glia in the retina of the rabbit. J Comp Neurol. 1985, 240(2), 128–42. doi: 10.1002/cne.902400203.

32. Zhang, J.; Wu, G.S.; Ishimoto, S.; Pararajasegaram, G.; Rao, N.A. Expression of major histocompatibility complex molecules in rodent retina. Immunohistochemical study. Invest Ophthalmol Vis Sci. 1997, 38(9), 1848–57.

33. Tsukamoto, Y.; Omi, N. Classification of Mouse Retinal Bipolar Cells: Type-Specific Connectivity with Special Reference to Rod-Driven AII Amacrine Pathways. Front Neuroanat. 2017, 11, 92. doi: 10.3389/fnana.2017.00092.

34. Bloomfield, S.A. Relationship between receptive and dendritic field size of amacrine cells in the rabbit retina. J Neurophysiol. 1992, 68(3), 711–25. doi: 10.1152/jn.1992.68.3.711.

35. Morin, L.P.; Studholme, K.M. Retinofugal projections in the mouse. J Comp Neurol. 2014, 522(16), 3733–53. doi: 10.1002/cne.23635.

36. Seabrook, T.A.; Burbridge, T.J. Crair MC, Huberman AD. Architecture, Function, and Assembly of the Mouse Visual System. Annu Rev Neurosci. 2017, 40, 499–538. doi: 10.1146/annurev-neuro-071714-033842.

37. Mannix, R.; Monuteaux, M.C.; Schutzman, S.A.; Meehan, W.P. 3rd; Nigrovic, L.E.; Neuman, M.I. Isolated skull fractures: trends in management in US pediatric emergency departments. Ann Emerg Med. 2013, 62(4), 327–31. doi: 10.1016/j.annemergmed.2013.02.027.

38. Colonna, M.; Butovsky, O. Microglia Function in the Central Nervous System During Health and Neurodegeneration. Annu Rev Immunol. 2017, 35, 441–468. doi: 10.1146/annurev-immunol-051116-052358.

39. Honig, M.G.; Del Mar, N.A.; Henderson, D.L.; O’Neal, D.; Doty, J.B.; Cox, R.; Li, C.; Perry, A.M.; Moore, B.M.; Reiner, A. Raloxifene Modulates Microglia and Rescues Visual Deficits and Pathology After Impact Traumatic Brain Injury. Front Neurosci. 2021, 15, 701317. doi: 10.3389/fnins.2021.701317.

40. Childs, C.; Barker, L.A.; Gage, A.M.; Loosemore, M. Investigating possible retinal biomarkers of head trauma in Olympic boxers using optical coherence tomography. Eye Brain. 2018, 10, 101–110. doi: 10.2147/EB.S183042.

41. Jin, N.; Gao, L.; Fan, X.; Xu, H. Friend or Foe? Resident Microglia vs Bone Marrow-Derived Microglia and Their Roles in the Retinal Degeneration. Mol Neurobiol. 2017, 54(6), 4094–4112. doi: 10.1007/s12035-016-9960-9.

42. Hernandez-Ontiveros, D.G.; Tajiri, N.; Acosta, S.; Giunta, B.; Tan, J.; Borlongan, C.V. Microglia activation as a biomarker for traumatic brain injury. Front Neurol. 2013, 4, 30. doi: 10.3389/fneur.2013.00030.

43. Kumar, S. Mechanisms mediating caspase activation in cell death. Cell Death Differ. 1999, 6(11), 1060–6. doi: 10.1038/sj.cdd.4400600.

44. Hengartner, M.O. The biochemistry of apoptosis. Nature. 2000, 407(6805), 770–6. doi: 10.1038/35037710. PMID: 11048727.

45. Knoblach, S.M.; Nikolaeva, M.; Huang, X.; Fan, L.; Krajewski, S.; Reed, J.C.; Faden, A.I. Multiple caspases are activated after traumatic brain injury: evidence for involvement in functional outcome. J Neurotrauma 2002, 19(10), 1155–70.

46. Glushakov, A.O.; Glushakova, O.Y.; Korol, T.Y.; Acosta, S.A.; Borlongan, C.V.; Valadka, A.B.; Hayes, R.L.; Glushakov, A.V. Chronic Upregulation of Cleaved-Caspase-3 Associated with Chronic Myelin Pathology and Microvascular Reorganization in the Thalamus after Traumatic Brain Injury in Rats. Int J Mol Sci. 2018, 19(10), 3151. doi: 10.3390/ijms19103151.

47. Boatright, K.M.; Salvesen, G.S. Caspase activation. Biochem Soc Symp. 2003, (0), 233–42. doi: 10.1042/bss0700233.

48. Boatright, K.M.; Salvesen, G.S. Mechanisms of caspase activation. Curr Opin Cell Biol. 2003, 15(6), 725–31. doi: 10.1016/j.ceb.2003.10.009.

49. Liu, Y.X.; Sun, H.; Guo, W.Y. Astrocyte polarization in glaucoma: a new opportunity. Neural Regen Res. 2022, 17(12), 2582–2588. doi: 10.4103/1673-5374.339470.

50. Gharagozloo, M.; Smith, M.D.; Jin, J.; Garton, T.; Taylor, M.; Chao, A.; Meyers, K.; Kornberg, M.D.; Zack, D.J.; Ohayon, J.; Calabresi, B.A.; Reich D.S.; Eberhart, C.G.; Pardo, C.A.; Kemper, C.; Whartenby, K.A.; Calabresi, P.A. Complement component 3 from astrocytes mediates retinal ganglion cell loss during neuroinflammation. Acta Neuropathol. 2021, 142(5), 899–915. doi: 10.1007/s00401-021-02366-4.

51. Xu, Q.A.; Boerkoel, P.; Hirsch-Reinshagen, V.; Mackenzie, I.R.; Hsiung, G.R.; Charm, G.; To, E.F.; Liu, A.Q.; Schwab, K.; Jiang, K.; Sarunic, M.; Beg, M.F.; Pham, W.; Cui, J.; To, E.; Lee, S.; Matsubara, J.A. Müller cell degeneration and microglial dysfunction in the Alzheimer’s retina. Acta Neuropathol Commun. 2022, 10(1), 145. doi: 10.1186/s40478-022-01448-y.

52. Wang, J.; Fox, M.A.; Povlishock, J.T. Diffuse traumatic axonal injury in the optic nerve does not elicit retinal ganglion cell loss. J Neuropathol Exp Neurol. 2013, 72(8), 768–81. doi: 10.1097/NEN.0b013e31829d8d9d.

53. Ma, J.; Zhang, K.; Wang, Z.; Chen, G. Progress of Research on Diffuse Axonal Injury after Traumatic Brain Injury. Neural Plast. 2016, 2016, 9746313. doi: 10.1155/2016/9746313.

54. Klimo, K.R.; Stern-Green, E.A.; Shelton, E.; Day, E.; Jordan, L.; Robich, M.; Racine, J.; McDaniel, C.E.; VanNasdale, D.A.; Yuhas, P.T. Structure and function of retinal ganglion cells in subjects with a history of repeated traumatic brain injury. Front Neurol. 2022, 13, 963587. doi: 10.3389/fneur.2022.963587.

55. Harper, M.M.; Boehme, N.; Dutca, L.M.; Anderson, M.G. The Retinal Ganglion Cell Response to Blast-Mediated Traumatic Brain Injury Is Genetic Background Dependent. Invest Ophthalmol Vis Sci. 2021, 62(7), 13. doi: 10.1167/iovs.62.7.13.

56. Villacampa, P.; Liyanage, S.E.; Klaska, I.P.; Cristante, E.; Menger, K.E.; Sampson, R.D.; Barlow, M.; Abelleira-Hervas, L.; Duran, Y.; Smith, A.J.; Ali, R.R.; Luhmann, U.F.O.; Bainbridge, J.W.B. Stabilization of myeloid-derived HIFs promotes vascular regeneration in retinal ischemia. Angiogenesis. 2020, 23(2), 83–90. doi: 10.1007/s10456-019-09681-1.

57. Kavanagh, E.; Rodhe, J.; Burguillos, M.A.; Venero, J.L.; Joseph, B. Regulation of caspase-3 processing by cIAP2 controls the switch between pro-inflammatory activation and cell death in microglia. Cell Death Dis. 2014, 5(12), e1565. doi: 10.1038/cddis.2014.514.

58. Todd, L.; Palazzo, I.; Suarez, L.; Liu, X.; Volkov, L.; Hoang, T.V.; Campbell, W.A.; Blackshaw, S.; Quan, N.; Fischer, A.J. Reactive microglia and IL1β/IL-1R1-signaling mediate neuroprotection in excitotoxin-damaged mouse retina. J Neuroinflammation. 2019, 16(1), 118. doi: 10.1186/s12974-019-1505-5.

59. Szabo, E.; Patko, E.; Vaczy, A.; Molitor, D.; Csutak, A.; Toth, G.; Reglodi, D.; Atlasz, T. Retinoprotective Effects of PACAP Eye Drops in Microbead-Induced Glaucoma Model in Rats. Int. J. Mol. Sci. 2021, 22, 8825. https://doi.org/10.3390/ijms22168825

60. Pellissier, L.P.; Hoek, R.M.; Vos, R.M.; Aartsen, W.M.; Klimczak, R.R.; Hoyng, S.A.; Flannery, J.G.; Wijnholds, J. Specific tools for targeting and expression in Müller glial cells. Mol Ther Methods Clin Dev. 2014, 1, 14009. doi: 10.1038/mtm.2014.9.

61. Zhang, C.; Guo, Y.; Slater, B.J.; Miller, N.R.; Bernstein, S.L. Axonal degeneration, regeneration and ganglion cell death in a rodent model of anterior ischemic optic neuropathy (rAION). Exp Eye Res. 2010, 91(2), 286–92. doi: 10.1016/j.exer.2010.05.021.

62. Durham, H.D. Demonstration of hyperphosphorylated neurofilaments in neuronal perikarya in vivo by microinjection of antibodies into cultured spinal neurons. J Neuropathol Exp Neurol. 1990, 49(6), 582–90. doi: 10.1097/00005072-199011000-00004.

63. Fricker, M.; Tolkovsky, A.M.; Borutaite, V.; Coleman, M.; Brown, G.C. Neuronal Cell Death. Physiological reviews 2018, 98(2), 813–880. doi: 10.1152/physrev.00011.2017

64. Burguillos, M.A.; Deierborg, T.; Kavanagh, E.; Persson, A.; Hajji, N.; Garcia-Quintanilla, A.; Cano, J.; Brundin, P.; Englund, E.; Venero, J. L.; Joseph, B. Caspase signalling controls microglia activation and neurotoxicity. Nature 2011, 472(7343), 319–324. doi: 10.1038/nature09788

